# ^1^H, ^13^C and ^15^N resonance assignment of the SARS-CoV-2 full-length nsp1 protein and its mutants reveals its unique secondary structure features in solution

**DOI:** 10.1101/2021.05.05.442725

**Authors:** Tatiana Agback, Francisco Dominguez, Ilya Frolov, Elena I. Frolova, Peter Agback

## Abstract

Structural characterization of the SARS-CoV-2 full length nsp1 protein will be an essential tool for developing new target-directed antiviral drugs against SARS-CoV-2 and for further understanding of intra- and intermolecular interactions of this protein. As a first step in the NMR studies of the protein, we report the ^1^H, ^13^C and ^15^N resonance backbone assignment as well as the Cβ of the apo form of the full-lengthSARS-CoV-2 nsp1 including folded domain together with the flaking N- and C-terminal intrinsically disordered fragments. The 19.8 kD protein was characterized by high-resolution NMR. Validation of assignment have been done by using two different mutants, H81P and K129E/D48E as well as by amino acid specific experiments. According to the obtained assignment, the secondary structure of the folded domain in solution was almost identical to its previously published X-ray structure, but some discrepancies have been detected. In the solution SARS-CoV-2 nsp1 exhibited disordered, flexible N- and C-termini with different dynamic characteristics. The short peptide in the beginning of the disordered C-terminal domain adopted two different conformations distinguishable on the NMR time scale. We propose that the disordered and folded nsp1 domains are not fully independent units but are rather involved in intramolecular interactions. Studies of the structure and dynamics of the SARS-CoV-2 mutant in solution are on-going and will provide important insights into the molecular mechanisms underlying these interactions.

## Introduction

Within the recent 1.5 years, the Severe Acute Respiratory Syndrome coronavirus 2 (SARS-CoV-2) has spread all over the world and devastated the economies of essentially all countries (1, 2). To date, more than one hundred million people have contracted the disease that led more than 3 M deaths (https://www.worldometers.info/coronavirus). Despite the enormous public health threat of this and previous CoV infections, no efficient therapeutic means have been developed against coronaviruses (CoV) before the COVID-19 pandemics. One of the major reasons for this was a lack of detailed knowledge of the mechanism of CoV replication and interaction with host cells.

SARS-CoV-2 is a member of the *Betacoronavirus* (β-CoV) genus along with other highly pathogenic respiratory viruses, such as SARS-CoV-1 and MERS-CoV (Middle Eastern respiratory syndrome virus). These viruses have similar genome and replication strategies but differ in their pathogenicity for humans. Similar to other β-CoVs, the SARS-CoV-2 genome (G RNA) is represented by a single-stranded RNA of positive polarity of ~30 kb in length (3–5). It mimics the structure of cellular mRNAs in that it has a Cap and a poly(A)-tail at the 5’ and 3’ termini, respectively. Upon delivery into the cells, the G RNA is directly translated into two very long polyproteins, which are encoded by the overlapping ORF1a and ORF1b. The latter polyproteins are enzymatically processed into individual nonstructural proteins nsp1-to-16 by the encoded protease activities. These nsps represent viral components of the replication complexes and are also involved in modification of the intracellular environment to promote efficient viral replication. As in the case of other β-CoVs, the SARS-CoV-2-specific nsp1 protein plays indispensable roles in these processes (6–8). First of all, it is a key player in downregulation of cellular translation and is a major β-CoV-specific virulence factor (6, 9, 10). It interacts with the 40S ribosomal subunit, blocks the RNA channel and inhibits initiation of translation of cellular, but not viral, RNA templates (6, 8, 11–17). SARS-CoV-1 and MERS nsp1 proteins are also indirectly involved in endonuclease degradation of cellular mRNAs; however, the mechanism of this function remains to be determined (18, 19). It is still unknown whether SARS-CoV-2 nsp1 can mediate degradation of cellular RNAs. Nsp1 of both SARS-CoV-1 and SARS-CoV-2 were also implicated in inhibition of nuclear-cytoplasmic traffic, albeit by different mechanisms (20, 21). The above activities appear to play critical roles in the downregulation of the innate immune response that can mount during SARS-CoV-2 infection, and thus, control the infection spread. Nsp1 proteins of β-CoVs also facilitate cell cycle arrest, which is clearly detectable during viral infection and nsp1 expression (9, 22). Importantly, the previous studies demonstrated that the deletion of nsp1 gene in the genome of other β-CoVs makes them nonviable (23). This strongly indicated the direct involvement of the latter protein in genomic RNA replication and/or synthesis of the subgenomic RNAs, which encode viral structural and accessory proteins. Interestingly, point mutations or small deletions in nsp1 can independently prevent either inhibition of cellular translation or viral replication (22–27).

Thus, the accumulated data suggest that nsp1 plays important roles in CoV replication and pathogenesis. It exhibits multiple activities and likely interacts with a variety of viral and cellular proteins and organelles. Understanding of the molecular mechanisms of these interactions is critical for development of live attenuated vaccines and therapeutic means against SARS-CoV-2 infection. Further dissection of multiple nsp1 functions in viral replication and pathogenesis requires the detailed knowledge of the dynamic structure of nsp1. To date, the data about the structure of β-CoV nsp1 remain very fragmented. This is a relatively small 19.8 kDa protein. It contains an N-terminal structured domain (~aa 10 to 124 in SARS-CoV2 nsp1), which was proposed to be critical for degradation of cellular mRNAs. The first 10 aa and the C-terminal fragment (aa125-180) in SARS-Cov-2 nsp1 are predicted to be intrinsically disordered. However, the last 26-aa-long peptide in this C-terminal fragment has been shown to fold into two short α-helixes upon binding to 40S ribosome subunit (7, 8, 17). The structure of folded N-terminal domain of SARS-CoV nsp1 has been determined by NMR (PDB:2HSX), and two X-ray structures of the folded domain of SARS-CoV-2 nsp1 have been recently published (PDB:7K7P and 7K3N) (28–30). The N-terminal nsp1 domains of both viruses have similar folds. The important difference was found to be the presence of an additional small β-strand (aa 95-97) in SARS-CoV-2-specific nsp1. No structure of the full-length proteins containing the disordered C-terminal domain is available for the nsp1 of any β-CoV nsp1.

The flexibility of the disordered C- and N-terminal fragments in the SARS-CoV-2 nsp1 and their interactions with the folded domain may play the critical role(s) in protein functions. Solution NMR is the method of choice for studying such flexible regions in proteins. Most of the currently available NMR approaches and protocols are focused on elucidating the structure of either fully folded proteins (FP), which complement the crystallographic data, or the intrinsically disordered protein (IDPs). Complete backbone and side chain resonance assignment of NMR spectra of large proteins, containing both folded and disordered domains, is still a challenge due to (a) a high degree of divergence in the conformational flexibility characteristics of disordered and folded domains and (b) the reduced frequency dispersion observed in the NMR spectra in the ^1^H dimension for disordered regions. Recently, a near-complete backbone resonance assignment of the SARS-CoV-2 nsp1 was reported (31). The latter protein was analysed in an acidic buffer, pH 6.5.

As a first step towards characterizing the structure and dynamics of the full-length SARS-CoV-2 nsp1 in neutral buffer by NMR spectroscopy, we herein report the almost complete ^1^H, ^13^C and ^15^N backbone and ^13^Cβ side chain assignment of the wild type protein and two of its mutants: a single mutant H81P and a double mutant K129E, D48E. This assignment has been evaluated by additionally observing selectively chosen type of amino acids (MUSIC-type of experiments). To overcome ambiguities in assignment in the crowded areas of the full-length SARS-CoV-2 nsp1 spectra, we additionally compared the assigned resonances with those corresponding to the single and double mutants of nsp1. Based on this data, the secondary structure of the full-length SARS-CoV-2 nsp1 in solution was derived and, additionally, the protein flexibility was evaluated. These data provide a structural basis for further understanding of intra- and intermolecular interactions of the SARS-CoV-2 nsp1.

## Material and Methods

### Preparation of ^15^N, ^13^C labeled SARS-CoV-2 nsp1 proteins

Nucleotide sequence of full length wt nsp1 was amplified by PCR from recombinant cDNA of SARS-CoV-2 Wuhan-Hu-1 strain (NC_045512.2) using primers:

NSP1-Dir CCACTGGTCTCAAGGTGGTATGGAGAGCCTTGTCCCTGG
NSP1-Rev CCACACTCGAGTTATTACCCTCCGTTAAGCTCACGC.

The PCR product was cloned into pE-SUMOpro-3 plasmid (LifeSensors Inc) between Bsa I and Xho I restriction sites. The synthetic gene blocks encoding mutant nsp1 were ordered from Integrated DNA Technologies and cloned into pE-SUMOpro-3 plasmid (LifeSensors Inc) between Eco RI and Xho I restriction sites. Plasmids encoding SUMO-nsp1 proteins were transformed into *E. coli* strain Rosetta2(DE3)pLacI (Novagen), and proteins were produced in the M9 media supplemented with 2 g/L [^15^N]NH_4_Cl (Cambridge Isotope Laboratories) and 3 g/L d-[^13^C_6_]glucose (Cambridge Isotope Laboratories). The expression was induced by 1 mM IPTG after cells reached the density of ~2 OD_600_. Then cells continued to grow at 37°C for 3-5 h. Freshly prepared or frozen cell pellets were lysed in Emulsiflex B15 (Avestin). The lysates were loaded on HisTrap HP column (GE Healthcare) and after extensive washing the recombinant proteins were eluted by imidazole gradient. Fractions containing SUMO-HVD nsp1 proteins were combined, and His-SUMO tag was cleaved with Upl1 protease. After cleavage, the proteins were diluted to contain 25 mM NaCl and further purified on Resource Q column (GE Healthcare). Size exclusion chromatography on a HiLoad Superdex 75 16/600 column (GE Healthcare) in NMR buffer was used as a final purification step. Fractions containing pure proteins were combined and concentrated.

The protein purities and identities were confirmed by SDS-PAGE and mass spectrometry (for unlabeled protein), respectively. Final proteins contained extra glycine at the N-terminus, which was required forSUMO cleavage. Protein concentrations were determined on 280 nm using extinction coefficients, which were determined by ProteinCalculator v3.4 (http://protcalc.sourceforge.net/).

### NMR samples preparation

All NMR experiments were performed in a buffer containing 20 mM HEPES pH 7.5, 100 mM KCl (with or without 2.5 mM MgCl_2_), 1 mM NaN_3_, 10 (v/v) % D_2_O and 0.1 mM DSS (4,4-dimethyl-4-silapentane-1-sulfonic acid) as an internal ^1^H chemical shift standard. The protein concentration were about 0.4 mM and spectra were acquired in a 5 mm tube (final volume of 0.45mL). ^13^C and ^15^N chemical shifts were referenced indirectly to the ^1^H standard using a conversion factor derived from the ratio of NMR frequencies(32).

To perform 3D ^13^C-HSQC type experiments, the nsp1 sample was lyophilized and dissolved in D_2_O in a volume of 0.45 mL. To confirm that the protein’s structure was not affected by lyophilization, sample was lyophilized again, dissolved in H_2_O and 1D ^1^H spectrum was compared to the original spectrum.

### NMR experiments

NMR experiments were acquired on Bruker Avance III spectrometers operating at 14.1 T, equipped with a cryo-enhanced QCI-P probe. The assignment of the backbone and side chains resonances was based on a set of 3D TROSY or HSQC triple resonance experiments from the Bruker library. The summary of the performed experiments and the key parameters are presented in Table 1. To increase resolution in the indirect dimensions and reduce acquisition time of the 3D experiments, the iterative non-uniform sampling protocol (NUS) (33) comprised TROSY-HNCO, TROSY-HNCA and TROSY-HN(CO)CA, TROSY-HN(CA)CO, TROSY-HN(CO)CACB, TROSY HNCACB, H(CC)(CO)NH, DIPSI2-^15^N HSQC, HCACO and MLEV17-^13^C HSQC experiments. NUS points sampling schedule applied to 3D experiments are listed in Table 1.

**Table 1.**
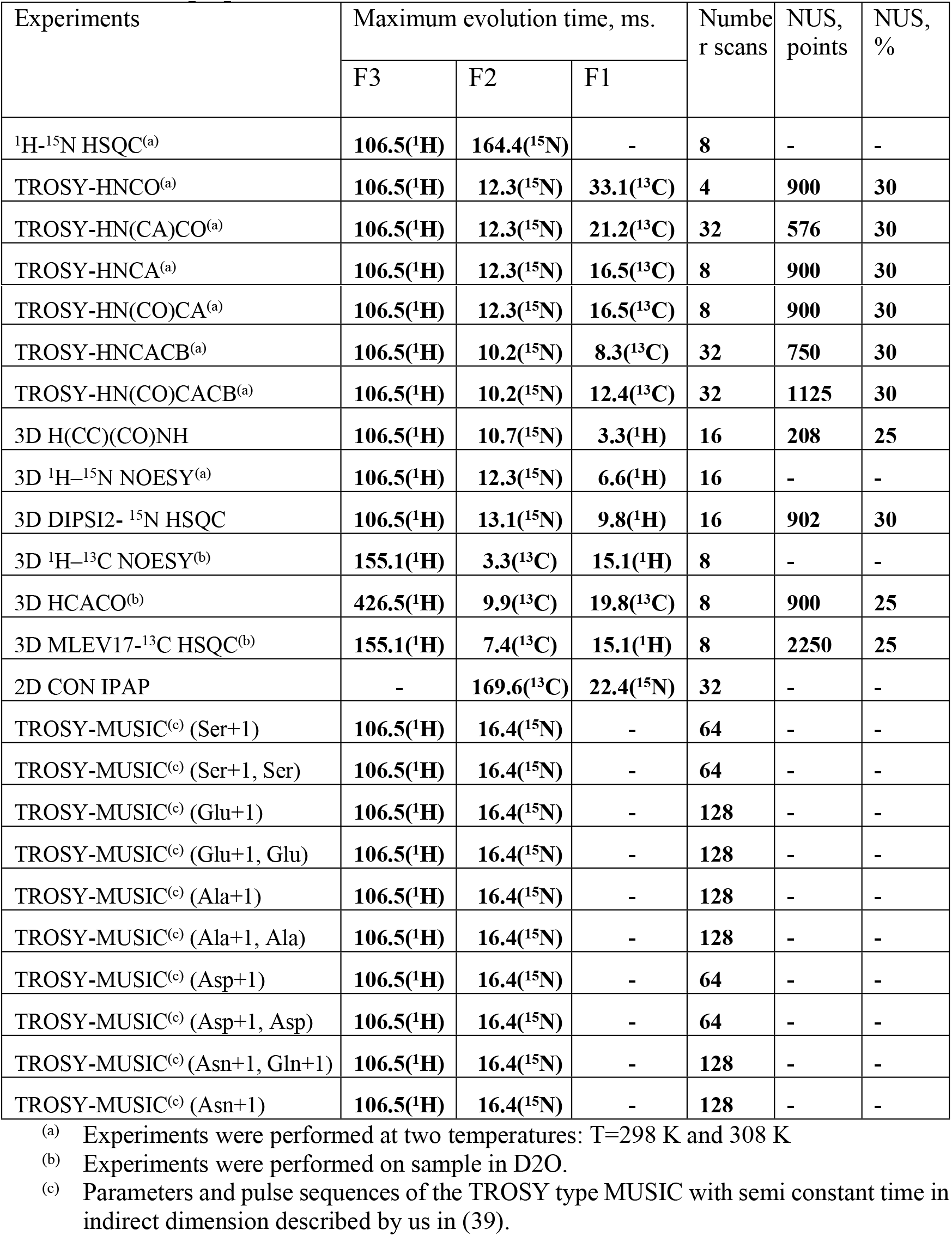
List of NMR experiments and the main parameters used to perform the sequence specific assignment of the backbone and sidechains of the full-length SARS-CoV-2 nsp1 protein.

To assign Hα proton resonances, additional ^1^H-^15^N-NOESY, ^1^H-^13^C-NOESY (34–36) data were collected.

To verify assignment of the aa located in the disordered fragment of SARS-CoV-2 nsp1, ^13^C observed CON experiment with IPAP scheme for virtual decoupling (37, 38) was used to correlate ^15^N with 13C’ resonances.

We also performed TROSY type MUSIC experiments with semi-constant time acquisition period in the indirect dimension to identify different type of aa such as Ser, Glu, Ala, Asp, Gln, Asn and their respective n+1 residues. Key parameters of the experiments used in this study are presented in Table 1. All TROSY-type MUSIC pulse sequences and the setting details have been fully described by elsewhere (39).

Due to the differences in the relaxation characteristics of the folded domain and the disordered fragments of SARS-CoV-2 nsp1, several experiments were performed at two temperatures: 298 K and 308 K.

Data were processed by Topspin 4.0.6 (Bruker) and assigned using CcpNmr Analysis 2.4.2

(40).

The chemical shifts of the full-length SARS-CoV-2 nsp1 were analyzed with TALOS+ software (41). As input for TALOS+ analysis, the experimentally derived chemical shifts of ^1^HN, ^15^N, ^13^Cα, ^13^Cβ, ^13^C’ and ^1^Hα nuclei for every aa were used. In case of absence of the chemical shifts, TALOS+ uses a database of sequences to predict the secondary structure (41).

The secondary X-ray based structures were extracted with the UCSF Chimera program(42) using the PDB entry: 7K7P (30).

In the text and figures, the standard nomenclature for amino acids of the carbon atoms was used, where ^13^Cα is the carbon next to the carbonyl group ^13^C’ and ^3^Cβ is the carbon next to ^13^Cα (43).

## Result and discussion

### Assignment protocol

The validity of the secondary structure analysis based on chemical shifts (CS) depends on the knowledge of CS of the ^1^HN, ^15^N, ^13^Cα, ^13^Cβ, ^13^C’ nuclei but also on ^1^Hα assigned resonances. Thus, we used the fully protonated ^15^N,^13^C-labelled SARS-CoV-2 nsp1. Additionally, this provides a foundation for validating inter β-strands interactions in the folded domain of the protein through observation of nOe contacts between ^1^HN-^1^HN or Hα - ^1^HN protons (44). Recently, it was reported (31) that the secondary structure of the full-length SARS-CoV-2 nsp1 protein (Fig. 1A) at pH 6.5 embraces one folded domain (aa 14-125) and two disordered chains, flanking the folded domain at the N-teminus (aa 1-13) and the C-terminus (aa 126-180). It is known that folded and intrinsically disordered proteins (IDP) entities, have differences in relaxation properties and, thus, require different optimal NMR experimental conditions. This also means that the positions of the amide ^1^H and ^15^N chemical shifts of the disordered fragments and some flexible parts of the folded domains strongly depend on the buffer conditions including pH and temperature. Keeping this in mind, we performed all NMR data collections at two temperature, 298 K and 308 K. Consequently, we achieved the maximum NMR performances for the folded domain at 308 K and for the disordered fragments at 298 K. We also performed all assignments at physiological buffer (pH 7.5), low salt, and in the absence or presence of 2.5 mM MgCl_2_. These conditions are optimal for future analysis of nsp1 interaction with 40S ribosome or RNA.

**Fig 1.**
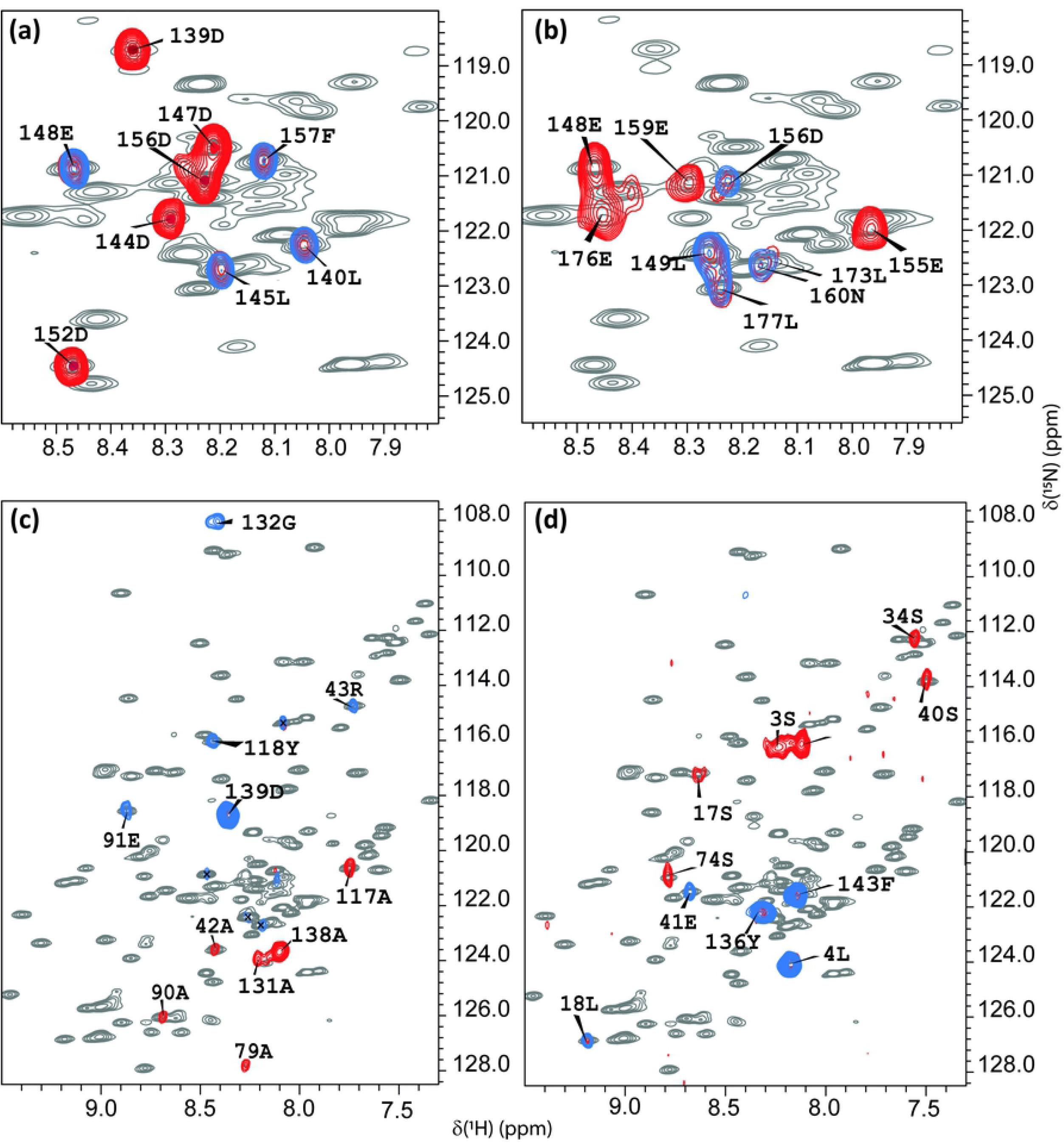
^1^H-^15^N HSQC spectra at T=308K of the full-length SARS-CoV-2 nsp1 protein with TROSY MUSIC experiments. Superpositions of the ^1^H-^15^N HSQC spectra at T=308K of the full-length SARS-CoV-2 nsp1 protein (grey) with TROSY MUSIC experiments performed at 308 K. (**a**) (blue) D + 1 and D (red) cross peaks; (**b**) (blue) E + 1 and E (red) cross peaks; (**c**) (blue) A + 1 and A (red) cross peaks; (**d**) (blue) S + 1 and S (red) cross peaks. The assignment of the observed cross peaks was done according to the aa sequence, which is shown in Fig. 2A.

As there are no programs readily available for performing automatic assignment of a protein containing both folded and disordered regions, we use the conventional manual assignment strategy based on experiments presented in Table 1. To achieve the best resolution in the 3D experiments in the indirect dimensions and to resolve resonances corresponding to the disordered part of protein, NMR experiments were mostly performed with the NUS option (33).

To facilitate the assignment procedure in this study, we chose the following strategy. First, selective type TROSY-MUSIC experiments on full length wild type ^15^N,^13^C labelled SARS-CoV-2 nsp1 at 308 K were performed. Fig. 1 presents the superposition of the ^1^H-^15^N HSQC of SARS-CoV-2 nsp1 protein (shown in grey) with TROSY-MUSIC spectra of selected D + 1, D, E + 1, A + 1, A, S + 1 and S cross peaks of aa. As we have described earlier(39), this type of experiments mostly benefits analysis of IDPs.

Indeed, spectra of the TROSY-MUSIC presented in Fig. 1 (a) and (b) show that all D + 1, D, E + 1, E cross peaks corresponding to the disordered fragments, but not for the folded domain, of SARS-CoV-2 nsp1 protein were observed and allowed their discrimination. TROSY-MUSIC with A + 1, A, S + 1 selection were showed almost all expected correlations for their respective types of aa in the sequence through the full length of the nsp1 protein. The exception was the amide protons that are involved in a slow conformational exchange, and thus, their cross peaks in ^1^H-^15^N HSQC spectrum were broadened below the detection limit.

These data were used to assign the resonances at 308 K. The ^1^H-^15^N-HSQC spectrum at 308K shows well-dispersed and narrow-line widths of the amide signals (Fig. 2 B, C). At this temperature, we have observed and assigned 158 aa, including prolines. Importantly, even at this higher temperature (308K), amino acids 125K, 124R, 123L and 122L shows two sets of amides ^1^HN - ^15^N cross peaks, which allowed us to conclude that the aa between the folded domain and the C-terminal disordered part of SARS-CoV-2 nsp1 protein adopt two distinguishable conformations detectable in the NMR time scale.

**Fig 2.**
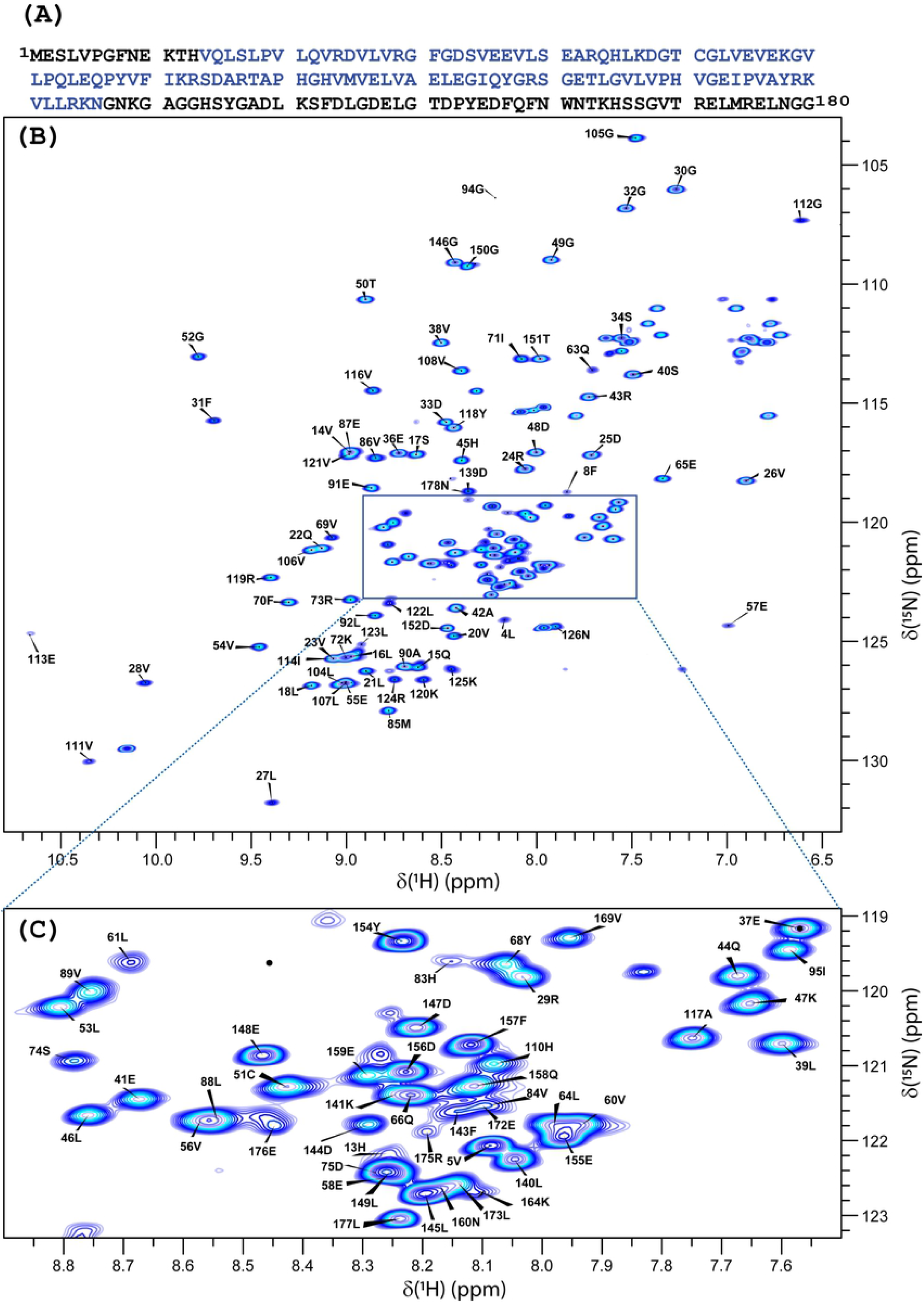
^1^H-^15^N HSQC spectra at T=308K of the full-length SARS-CoV-2 nsp1 protein with assignments. (**A**) Amino acid sequence of the full-length SARS-CoV-2 nsp1 protein. The folded domain is indicated in blue. (B) ^1^H-^15^N HSQC spectrum at 308K with its extended crowded part (**C**) of the full-length ^15^N, ^13^C-labelled SARS-CoV-2 nsp1 protein at 0.4 mM concentration in the buffer containing 20 mM HEPES pH 7.5, 100 mM KCl, 2.5 mM MgCl_2_, 1 mM TCEP, 1 mM NaN_3_, 10 (v/v) % D_2_O and 0.1 mM DSS at 308 K. The chemical shift assignment of NH backbone is shown by the number and symbols corresponding to the sequence (**A**). The assignment is presented only for the aa, whose cross peaks were observed at T 308K.

Next, we wanted to examine the broadening of some of the NH backbone resonances below the detection limit at 308K. To perform this analysis and additionally validate resonances ambiguously assigned due to its crowdedness in the spectra, we used NMR data of two SARS-CoV-2 nsp1 protein mutants. The selection of mutants was based on the following criteria: (1) the replacement of chosen aa should not lead to any strong conformational transition in the protein and (2) preferably large chemical shift perturbation (CSP) should be expected in the place of the exchanged aa. We have analysed NMR data of 2 mutants. In the first, a single histidine in position 81 was replaced by a proline (H81P). According to the X-ray structure (PDB: 7K7P), the aromatic ring of H81 is located in a flexible loop and turned towards the solvent. Thus, its replacement should not lead to any significant change in the architecture of the secondary structure of SARS-CoV-2 nsp1 protein. Moreover, the replacement by proline was expected to induce a large perturbation of chemical shifts (CS) of the nuclei located in its proximity due to its unique structure and possibly enhance the stability of the loop. As it is shown in Fig. 3d, the CSP observed in ^1^H-^15^N HSQC spectra of wt nsp1 vs the H81P mutant (red bars) are evident. As expected, the most significant CSPs were observed between aa 75 and 85. Noteworthy, the aa of the N-termini (10–17) and at the beginning of the C-terminal disordered fragment (120-127aa) are affected as well. This finding led us to the conclusion that aa corresponding to those three regions are in close proximity.

**Fig 3.**
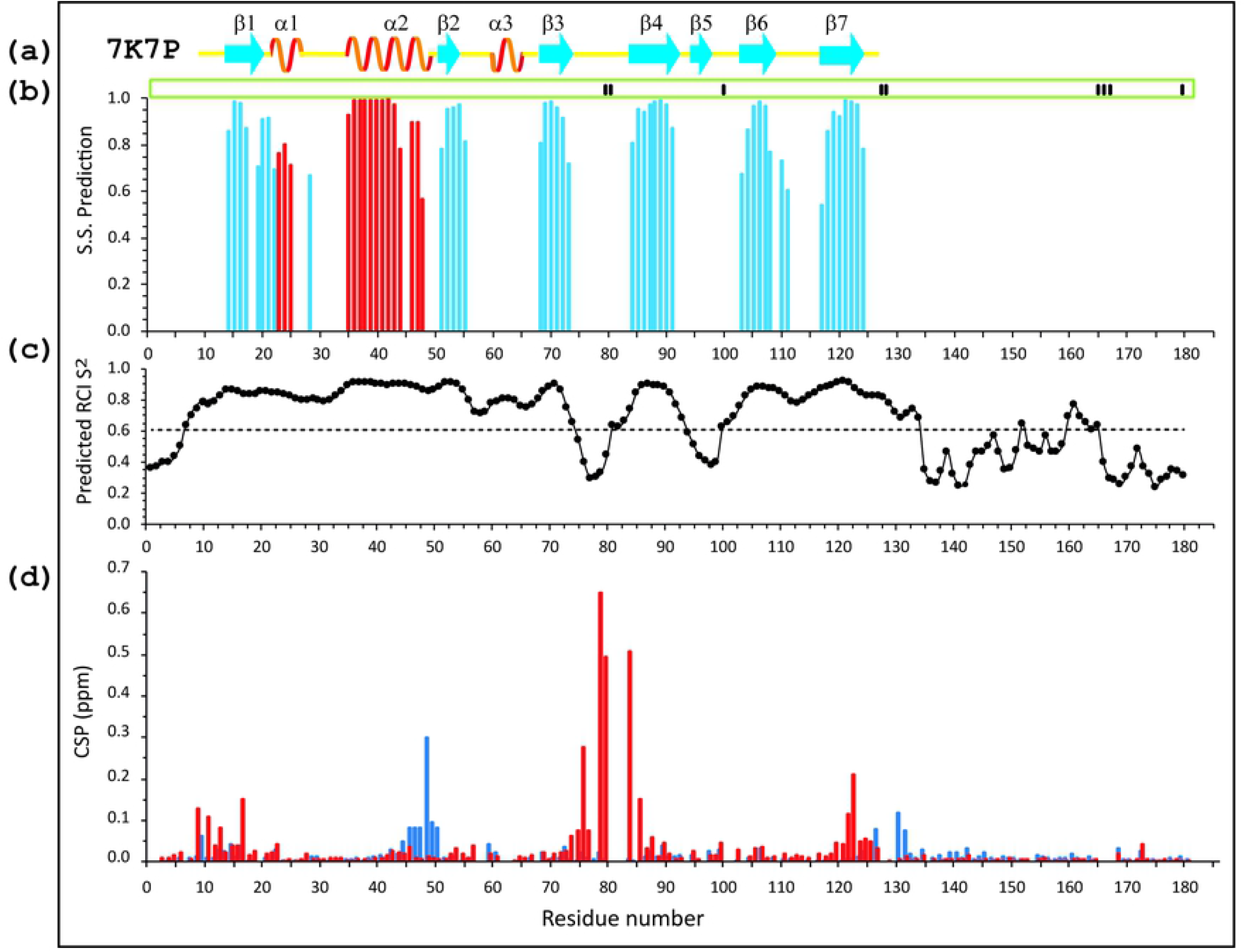
Secondary structure and order parameters of the full-length SARS-CoV-2 protein. **(a)** The secondary structure derived from the X-ray data of the folded domain of SARS-CoV-2 nsp1 (10-124 aa, PDB: 7K7P) according to UCSF Chimera (42). The yellow lines show loop segments of the protein. (**b**) Index of secondary structure prediction (S.S. Prediction) (*red* and *blue* bars indicate *α*-helix and *β*-strands, respectively. (**c**) Model-free (45) order parameter, S^2^, using random coil index (RCI)(46) extracted by TALOS+ (41). (**d**) Chemical shift deviations of Δ^1^H and Δ^15^N nuclei (ppm), obtained as distances 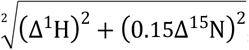, between wild type SARS-CoV-2 nsp1 protein and nsp1(H81P) mutant (red) or nsp1(K129E, D48E) mutant (blue). AAs, which did not contribute to the TALOS+ analysis are shown by short black bars in green box on top of panel (**b**).

In the second mutant, we introduced two aa substitutions: Lys in position 129 with positive charged side chain and Asp in position 48 with negative charged side chain were replaced by Asp with negative charged side chain (K129E and D48E). First mutation is in the beginning of the disordered region of SARS-CoV-2 nsp1. The second mutation, according to the X-ray structure (PDB: 7K7P), is located at the end of the α-helix. In Fig. 3d, the CSPs observed in ^1^H-^15^N HSQC spectra of wt nsp1 vs the double mutant is presented by blue bars. The significant CSPs were observed for aa 44-51 and 125-132.

The strategy applied in this study of inducing the CSP of nuclei in local area of protein through deliberate mutation of aa structurally insensitive was very valuable and helped us to resolve some ambiguities in the backbone and side chain assignments. More detailed characterisation of the SARS-CoV-2 nsp1 mutants will be published elsewhere.

The resulting assignment of the full length of SARS-CoV-2 nsp1 was as following. For the folded domain (aa 14-125) we assigned 95% ^1^HN and 95% ^15^N including prolines, 96% of ^13^Cα, 88% of ^13^Cβ, 94% of ^13^C’and 83% of all H^α^. For the two disordered fragments (aa 1-13 and aa 126-180) we assigned 91% ^1^HN, 91% ^15^N, 94% of ^13^Cα, 93% of ^13^Cβ, 94% of ^13^C’ and 76% of all H^α^.

Comparison of NMR spectra acquired in different conditions in this and published (31) work revealed only small changes for folded, dynamically stable nsp1 domain. However, the resonances corresponding to the loops and the disordered regions of the protein were strongly affected by changes in pH or temperature. All ^1^H, ^15^N and ^13^C chemical shifts of the full-length of SARS-CoV-2 nsp1 protein at pH 7.5 and at two temperatures, 298K and 308K, have been deposited in BioMagResBank (http://www.bmrb.wisc.edu) under the accession 50915. The assignment of the single mutant nsp1(H81P) and double mutant nsp1(K129E, D48E) can be provided upon reasonable request.

### Secondary structure of the SARS-CoV-2 nsp1 protein

The full length SARS-CoV-2 nsp1 protein chemical shifts were analyzed with TALOS+ (41), and the data are presented in Figs. 3b and c. The analysis of the secondary structures of the folded domain of the SARS-CoV-2 nsp1, which was derived from the NMR data, and the previously determined crystal structure (7K7P) shows that they are almost identical, and this has validated our resonance assignment (Fig. 3b and a). Nevertheless, a few important inconsistencies were identified. The crystal structure of SARS-CoV-2 nsp1 folded domain (28–30) revealed the presence of an additional short β5-strand (aa 95-97), which is not found in the structure of SARS-CoV nsp1 determined by NMR (PDB: 2HSX). In our study of the full-length nsp1, the presence of β5-strand folded between aa 95 and 97 could not be confirmed. Moreover, according to the predicted order parameter [S^2^ by TALOS+ (Fig. 3C)], the segment between aa 92-103 is dynamic. This prediction is in agreement with our finding that amide protons between residues I95 and G98 were not observed at 308K and 298K, suggesting their involvement in multiple conformational exchange and exposure to the solvent. We additionally performed an analysis of the 3D ^1^H-^15^N NOESY spectrum to determine dipole-dipole contacts of NH-NH and NH-Hα protons, which allows to detect hydrogen bonds between two β-strands (44). The β-sheet formed by strands β4 and β3, according to the X-ray structure, was confirmed by observing NH-NH and NH-Hα NOEs between those strands, but not between the β4 and β5 strands. These data contradict the X-ray results, which suggest low mobility of the β5 strand due to the additional hydrogen bonds between the β4 and β5 strands.

Subtle differences between the X-ray and NMR secondary structures were also noticed for the strands β1, β2, β6 and β7. According to our NMR data, in solution, these strands are extended by one or two aa at their C-termini. Furthermore, the α-helix 2 in solution is one aa shorter than in the X-ray structure. Our data also predicted that the α-helix has a break at aa H45. Another difference is that the previously identified short α-helix 3 is not predicted by the NMR data. Instead, it suggests the presence of a long, disordered loop between aa 55 and 67, which, according to TALOS+ prediction, has restricted mobility (Fig. 3b, c). Importantly, this region of SARS-CoV-2 nsp1 sequence is well characterized by NMR through CS as well as NOE of NH-NH, NH - NH-Hα protons contacts. This led us to conclusion that these discrepancies between NMR and X-ray secondary structure predictions are likely result from the crystallisation conditions.

Two loop regions in the folded domain of SARS-CoV-2, (aa Q27-S34 and G112-A117) are in agreement with the X-ray structure and have restricted mobility according to the higher order S^2^ parameter predicted by TALOS+ (Fig 3c).

Two more dynamic regions in the solution structure of the folded domain of SARS-CoV-2 were identified based on the S2 order parameter predicted by TALOS+: S74-H83 and L92-E102 (Fig 3b, c). This prediction is in line with the lack of peaks or broadened ^15^N/ ^1^HN cross peaks, even at T298K in the ^1^H-^15^N NMR spectra. We did not observe resonances for H81 and G82 in the first region and for S100 and G101 in the second one. It can be explained either by broadening of the cross peaks below detection limit or, which is more likely, by the involvement of these regions in slow conformational exchange.

The N- and C-termini, comprising aa M1-N9 and N124-G180, respectively, were identified by CSI as fully unstructured, but having different predicted order parameters (S^2^) through the sequences. Dynamic regions with an order parameter S^2^ below 0.6 were predicted for M1-F8, S135-Q158 and H165-G180. The increase in dynamic behaviour of those aa correlated with the changes in the intensities of the ^15^N/ ^1^HN backbone cross peaks in the ^1^H-^15^N NMR spectra of nsp1. These cross peaks have higher intensity compared to cross peaks belonging to the folded less dynamic aa. Based on these data, we propose that in the full-length SARS-CoV-2 nsp1, the folded and disordered parts of the protein behave not as fully independent units but are rather involved in intramolecular interactions.

In conclusion, the near complete ^15^N/^13^C/^1^H backbone resonance and part of side chain assignment of the full-length SARS-CoV-2 nsp1 at pH 7.5 and physiological salt concentration has been performed. Validation of assignment have been done by using two different nsp1 mutants as well as MUSIC type experiments. Assignment revealed that the secondary structure of the rigid folded domain is almost identical to that determined by X-ray. However, the existence of the short β-strand (aa 95 to 97), which is considered to be the significant structural difference between SARS-CoV-1 and SARS-CoV-2 nsp1 proteins, was not confirmed. In solution, SARS-CoV-2 nsp1exhibits disordered, flexible N- and C-termini, having different dynamics. The short peptide in the beginning of the C-terminal disordered fragment adopts two conformations. We propose that there are intramolecular interactions between the disordered and folded nsp1 domains. Studies of the structure and dynamics of the SARS-CoV-2 mutant in solution are on-going and will provide important insights on the molecular bases underlying these interactions.

## Acknowledgment

This work was supported by Swedish Foundation for Strategic Research grant ITM17-0218 to P.A., Public Health Service grant R21AI146969 to I.F. and UAB Research Acceleration Funds to E.F. and I.F. We thank Nikita Shiliaev for technical assistance.

## Conflicts of interest

The authors have no conflict of interest to declare.

## Notes

### Competing Interest Statement

The authors have declared no competing interest.

